# Multi-scale models reveal hypertrophic cardiomyopathy MYH7 G256E mutation drives hypercontractility and elevated mitochondrial respiration

**DOI:** 10.1101/2023.06.08.544276

**Authors:** Soah Lee, Alison S. Vander Roest, Cheavar A. Blair, Kerry Kao, Samantha B. Bremner, Matthew C Childers, Divya Pathak, Paul Heinrich, Daniel Lee, Orlando Chirikian, Saffie Mohran, Brock Roberts, Jacqueline E. Smith, James W. Jahng, David T. Paik, Joseph C. Wu, Ruwanthi N. Gunawardane, James A. Spudich, Kathleen Ruppel, David Mack, Beth L. Pruitt, Michael Regnier, Sean M. Wu, Daniel Bernstein

**Author notes:** **Corresponding author**: Daniel Bernstein, 240 Pasteur Dr Rm 2262, Palo Alto, California, 94304-1049. Equal Contribution (Soah Lee, Alison S. Vander Roest, Cheavar A. Blair, Kerry Kao, Samantha B. Bremner, Matthew C Childers, Divya Pathak).

## Abstract

**Rationale:** Over 200 mutations in the sarcomeric protein β-myosin heavy chain (MYH7) have been linked to hypertrophic cardiomyopathy (HCM). However, different mutations in MYH7 lead to variable penetrance and clinical severity, and alter myosin function to varying degrees, making it difficult to determine genotype-phenotype relationships, especially when caused by rare gene variants such as the G256E mutation.

**Objective:** This study aims to determine the effects of low penetrant MYH7 G256E mutation on myosin function. We hypothesize that the G256E mutation would alter myosin function, precipitating compensatory responses in cellular functions.

**Methods:** We developed a collaborative pipeline to characterize myosin function at multiple scales (protein to myofibril to cell to tissue). We also used our previously published data on other mutations to compare the degree to which myosin function was altered.

**Results:** At the protein level, the G256E mutation disrupts the transducer region of the S1 head and reduces the fraction of myosin in the folded-back state by 50.9%, suggesting more myosins available for contraction. Myofibrils isolated from hiPSC-CMs CRISPR-edited with G256E (MYH7^WT/G256E^) generated greater tension, had faster tension development and slower early phase relaxation, suggesting altered myosin-actin crossbridge cycling kinetics. This hypercontractile phenotype persisted in single-cell hiPSC-CMs and engineered heart tissues. Single-cell transcriptomic and metabolic profiling demonstrated upregulation of mitochondrial genes and increased mitochondrial respiration, suggesting altered bioenergetics as an early feature of HCM.

**Conclusions:** MYH7 G256E mutation causes structural instability in the transducer region, leading to hypercontractility across scales, perhaps from increased myosin recruitment and altered crossbridge cycling. Hypercontractile function of the mutant myosin was accompanied by increased mitochondrial respiration, while cellular hypertrophy was modest in the physiological stiffness environment. We believe that this multi-scale platform will be useful to elucidate genotype-phenotype relationships underlying other genetic cardiovascular diseases.

## INTRODUCTION

Hypertrophic cardiomyopathy (HCM), is the most common genetic heart disease affecting as many as 1 in 200 individuals. Complications of HCM include hypertrophy-related left ventricular outflow obstruction, heart failure, arrhythmia, and sudden cardiac death^1, 2^. HCM generally manifests with cardiac hypercontractility, cardiac hypertrophy, myocyte/myofibril disarray, and interstitial fibrosis^3^. Altered metabolism and mitochondrial function have more recently been described, even in the earlier phase of this disease^4^. About 60% of HCM cases are caused by familial inheritance of a single autosomal dominant mutation within a gene encoding for a sarcomere protein. In particular, the MYH7 gene encoding for β-myosin heavy chain (MHC) is reported to contain more than 200 mutations implicated in HCM, making it the second most common hotspot for HCM mutations^3, 5^. The majority of these mutations are found in the myosin head, where ATP and actin binding sites are located.

MYH7 mutations have variable effects and cause mutation-specific perturbation in myosin function. For example, earlier work from our group demonstrated that the H251N and D239N mutations, which are known to result in early-onset and severe HCM phenotypes in patients^6^, increase actin gliding velocity, ATPase activity and the fraction of active open-headed myosin compared to wild-type myosin, contributing to the observed hypercontractile phenotype^7, 8^ (**Table S1**). On the other hand, the R403Q mutation, commonly known as an adult-onset disease-causing mutation, has less severe effects *in vitro*, and its effects on myosin function have been debated because some have shown a gain-of-function^9–11^ (**Table S1**), whereas others have shown a loss-of-function^12, 13^.

This discrepancy likely resulted from different model systems used in previous studies. Reports on the effects of specific MYH7 mutations such as R403Q and P710R were not in agreement across scales (e.g. single-headed myosin^12, 14^, long-tailed myosin^9, 10, 14^, myofibril^13^, single cells^9, 14^, and tissues^15^). Moreover, previous studies have used various species as models including human, rabbit, and mouse, and examined the effects of MYH7 mutations on myosin function at only one or two scales^7, 8, 13, 14, 16–33^, perhaps leading to the observed variability in elucidating genotype-phenotype relationships. We propose that it is imperative to examine the effect of any mutation on myosin function across multiple scales and using a human-based system.

To address this, we assembled a multi-disciplinary collaborative team with expertise ranging from individual molecules to sub-cellular structures to whole cells to engineered micro-tissues to investigate the effects of HCM-specific MYH7 gene mutations. Among many MYH7 mutations, those that are rare or with insufficient or highly variable clinical data require comprehensive study using human-based multi-scale models to investigate mutation-specific pathogenic mechanisms. Among these is the MYH7 G256E mutation, which requires further genotype-phenotype correlation study since there is disparity between clinical phenotype data and structural prediction of the effects of the mutation on the myosin molecule. The G256E mutation is clinically classified as a pathogenic mutation and structurally predicted to cause a severe HCM phenotype due to the charge-changing nature of the mutation in the essential functional domain of myosin. However, one large kindred study suggested a benign nature of the G256E mutation. In this kindred (245 family members), 39 individuals (34 adults, 5 children) inherited the disease allele among 245 family members, and the G256E mutation showed low penetrance in both adults (19/34, 56%) and children (3/5, 60%). Only one instance of sudden cardiac death was described out of 245 family members despite pronounced cardiac hypertrophy^34^. This has led the G256E mutation to be classified by some as “benign” or pathogenic but associated with a lower relative risk of sudden cardiac death. However, more recent studies suggest that patients with previously labeled “benign” HCM mutations can experience serious adverse clinical outcomes^35^. Given the degree of uncertainty of the clinical significance of the G256E mutation, we selected this mutation to examine the genotype-phenotype correlation using our multi-scale models. We hypothesize that the G256E mutation would alter myosin function, precipitating compensatory responses in cellular functions.

Here we demonstrate the power of a multi-disciplinary collaborative team and a human protein/cell-based multi-scale system to comprehensively investigate the genotype-phenotype correlation. Using a CRISPR/Cas9 gene-edited hiPSC-CM model system and purified protein assays, we demonstrate a systematic workflow to reveal how the MYH7 G256E mutation affects myosin function, and the pathogenesis of HCM (**Figure 1**). Specifically, we (1) identified the angstrom level structural consequences of a single amino acid change using molecular dynamics simulations to study altered protein conformation^36–38^, (2) characterized how these protein structural changes affect myosin/actin interactions and the function of isolated myofibrils^20^, (3) identified how the changes in myosin function induce biomechanical and transcriptomic alterations in single hiPSC-CMs^39^, and finally (4) characterized the resulting impact at the tissue level using human engineered heart tissues (EHTs)^40, 41^.

**Figure 1.**
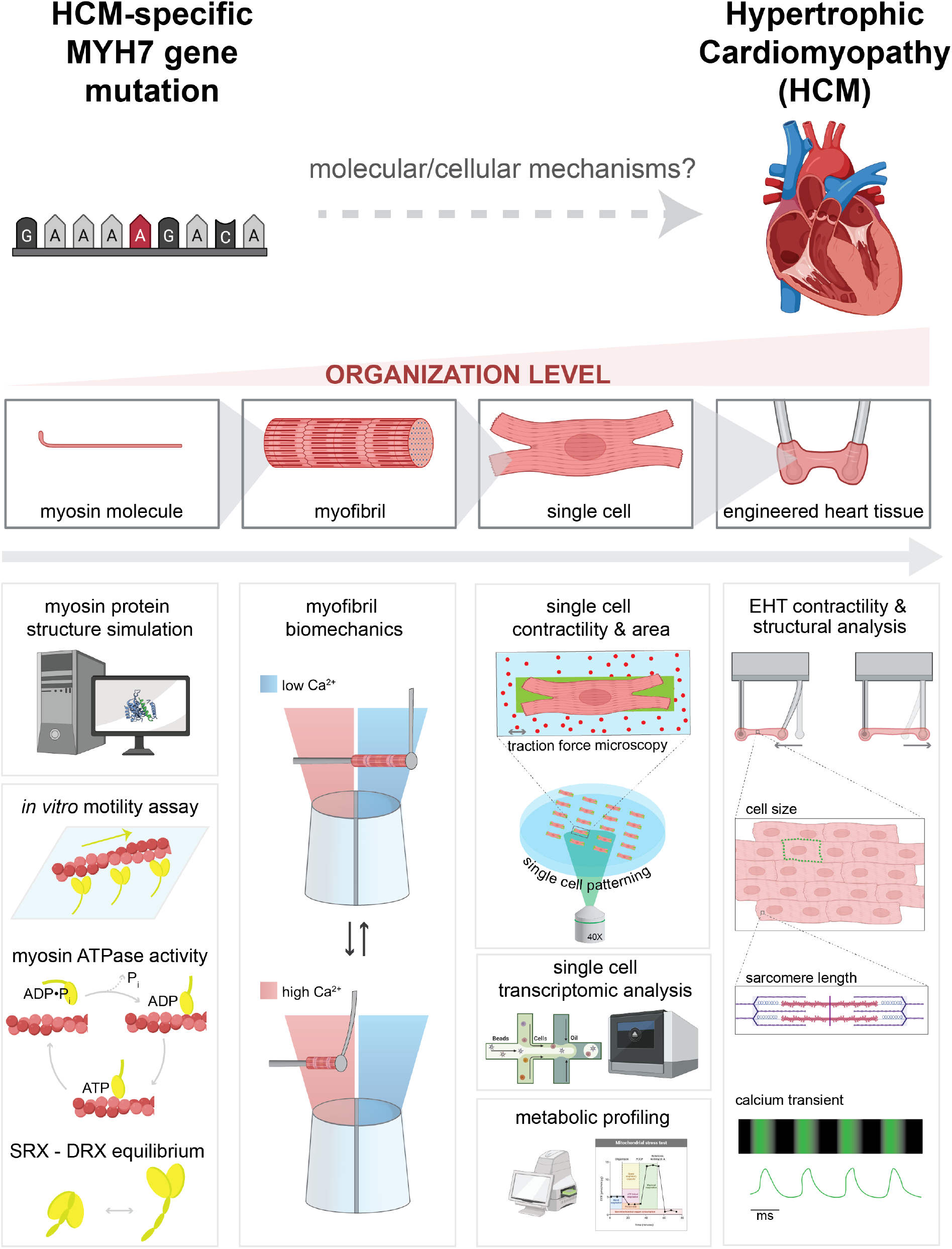
Schematic of multi-length scale examination of MYH7 mutation-driven HCM pathogenesis.

## METHODS

Full, extended Materials and Methods can be found in the Supplementary Information.

## RESULTS

### Molecular dynamics simulations

MYH7 G256E mutation is modeled to destabilize the myosin transducer region and alter structure near the ATP binding pocket.

G256 is located in a β-bulge hairpin turn that connects strands β_6_ and β_7_ of the myosin transducer region. The mutation site borders the myosin mesa and is sandwiched between an ɑ-helix of the N-terminal domain, loop 1, and the HO linker loop (**Figure 2a**). To predict mutation-driven conformational changes that result in altered myosin function, we used molecular dynamics (MD) simulations and modeled time-dependent structural changes of wild-type (WT) and G256E mutant (G256E) myosin at the atomic scale. In WT MD simulations, the β-bulge, and the transducer region in general, maintained a crystallographic-like conformation (**Figure 2a**). In the G256E simulation, however, replacement of the small and flexible Gly with the bulky and charged Glu led to a series of conformational changes. First, the backbone dihedral angles of the β-bulge were altered to accommodate the Glu. Second, the Glu side chain extended toward the N-terminal domain and formed an enduring salt bridge with R169 (**Figure 2b**). Formation of the E256-R169 salt bridge placed structural constraints on the transducer region, leading to statistically significant differences in side-chain to side-chain interactions among residues in this region including 4 interactions that were not sampled in the WT simulations (**Figure 2b,c**). In addition, the E256-R169 salt bridge placed increased strain on the transducer β-strands and led to statistically significant changes among backbone to backbone hydrogen bonds among β_5_, β_6_, and β_7_ (**Figure 2b,d**). Four of these backbone hydrogen bonds were present less often in the G256E simulations, which led to a decrease in structural stability of the transducer region and an increase in conformational heterogeneity among residues near the ATP binding pocket.

**Figure 2.**
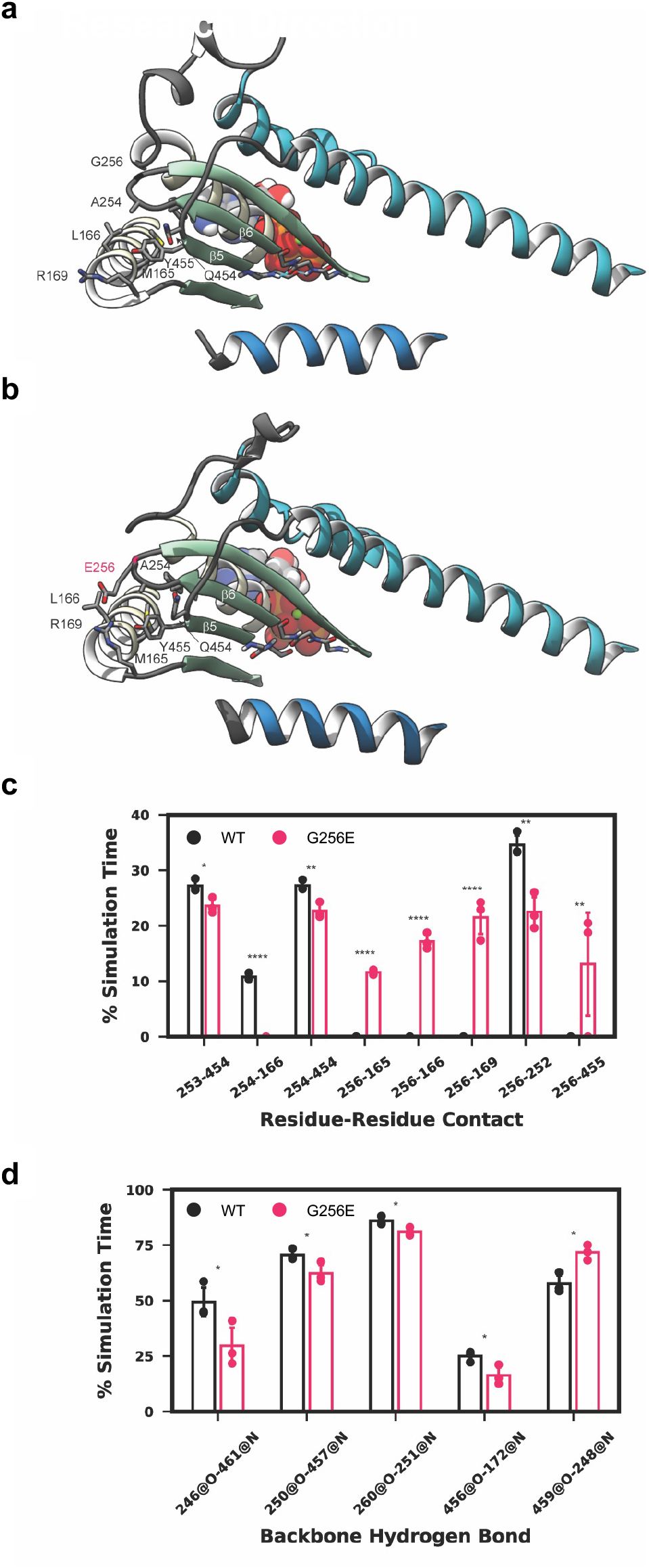
MYH7 G256E mutation disrupts the transducer region of myosin. The G256E mutation resulted in structural rearrangement of the transducer region of myosin, depicted in structural snapshots taken from the WT **(a)** and G256E **(b)** MD simulations of M.ATP state myosin. Residues in the transducer region are represented as ribbons, ATP Mg is represented as spheres. The sidechain and backbone atoms of residues with distinct atomic interactions are shown. **(c)** G256E resulted in a shift in residue-residue side chain contacts within the vicinity of the mutation site. Error bars are s.d. **(d)** G256E also resulted in a shift in backbone-backbone hydrogen bonding patterns among residues in the central β-sheet. Hydrogen bonds between strands β5 and β6 near the ATP pocket were formed less frequently in the G256E simulations. Data are presented as mean ± standard deviation (SD). Statistical significance was determined by a student’s t-test. *p ≦ 0.05, **p ≦ 0.01, ****p≦ 0.0001.

### Single molecule biomechanics

MYH7 G256E mutation reduces actin gliding velocity and alters the folded-back state of myosin.

To experimentally determine the effects of the G256E mutation on myosin motor function, we performed molecular assays with purified β-cardiac myosin constructs. Actin-activated ATPase, *in vitro* motility, and single nucleotide turnover assays were used to evaluate the impact of the G256E mutation on myosin kinetics and its interactions with actin. β-myosin heavy chain, the mechanoenzyme that drives ventricular contraction, forms a dimeric structure composed of two globular head/motor domains (S1) joined via an ɑ-helical coiled coil tail domain. Given the location of the G256E mutation in the head domain, we first used the single-headed short S1 form of β-cardiac myosin (sS1) to examine actin-myosin interactions^42, 43^. We found that the motility of actin filaments on the sS1-mutant-myosin-coated surface was reduced by 20% compared to the WT control (**Figure 3a**). However, the actin-activated ATPase rate (k_cat_) of the single myosin head domain (sS1) was not significantly altered by the G256E mutation, suggesting steady-state ATPase activity itself is not disrupted (**Figure 3b**). Next, we used double-headed β-cardiac myosin constructs (short-tailed form, 2hep and long-tailed form, 25hep) to determine the effect of the mutation on myosin intramolecular interactions. Myosin has been shown to regulate its activity by shifting the equilibrium between two conformation states^8, 10, 14, 44^: 1) a so-called super-relaxed state (SRX) that turns over ATP at very low rate and is thought to correlate to a structural state characterized by the S1 heads folding back onto the proximal tail (i.e. the interacting heads motif), and 2) a disordered-relaxed state (DRX) with a ∼10-fold higher ATPase rate that is characterized by open heads capable of attaching to actin and force generating. We found that in the case of WT myosin, actin-activated ATPase rates (k_cat_) for the long-tailed 25hep myosin construct are ∼40% lower than those of the short-tailed 2hep myosin construct that lacks the ability to form a folded-back state (**Figure 3c, Fig. S1**). However, the actin-activated ATPase rate of the long-tailed G256E mutant construct drops by only ∼20% compared to its short-tailed 2hep construct (G256E 2hep) (**Figure 3d, Fig. S1**). Since the only difference between the short- and long-tailed myosin constructs is the ability of the long-tailed construct to form a folded-back, sequestered state (**Figure 3b-d, Fig. S1**), this indicates that the smaller decrease in ATPase rate in the G256E mutant construct is likely due to disruption of the folded-back state (SRX), resulting in more myosin heads in an open state available to interact with actin. To further assess the balance between the folded-back “off” state (SRX) and open “on” state (DRX), a single turnover assay was performed using fluorescently labeled ATP. We found that the long-tailed 25hep construct containing the G256E mutation had a significantly lower proportion of myosin heads in the SRX state compared to WT 25hep myosin (p<0.0001) (**Figure 3e, Fig. S2;** WT 25hep= 54± 8.8% (mean± SD); G256E 25hep= 27.5± 10%), which indicates that G256E disrupts the folded-back state to increase myosin head availability for interacting with actin. To further validate that this increased SRX is tail-dependent, we assayed single-turnover ATP turnover kinetics for the 2-hep constructs, and we did not observe any significant change in SRX percentage for G256E compared to WT 2hep (WT= 19.7± 8.9% (mean ± SD); G256E= 22.6± 7.9%) (**Figure 3e, Fig. S2**). In sum, the G256E mutation led to a significant reduction in actin gliding velocity and the ability to form the folded-back state. The reduced ability to form the folded-back state may result in hypercontractility while the reduced actin gliding velocity may result from slower myosin-actin cycling, which could also result in hypercontractility under load. Thus, we next investigated the effects of the mutation at a higher organizational level in human myofibrils to understand the aggregate molecular effects of ensembles of motors on the ability to generate force.

**Figure 3.**
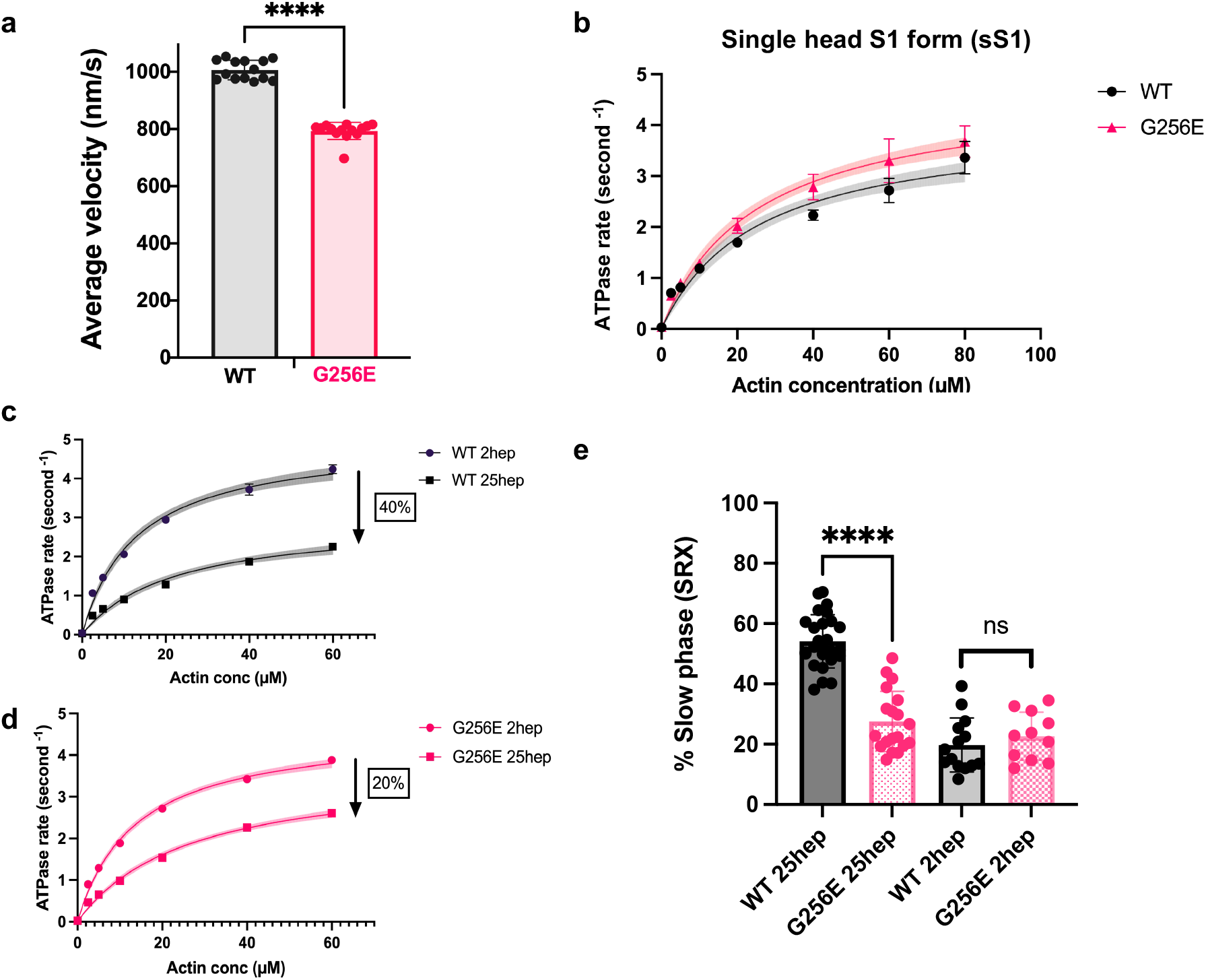
Measurements of myosin molecular activity revealed that G256E myosin exhibits a significant reduction in actin gliding velocity and super relaxed state (SRX) fraction. **(a)** Mean actin gliding velocities for wild-type (WT) and G256E short S1 (sS1) construct of β-myosin heavy chain. Mean actin gliding velocity shown (MVEL20) is the mean velocity of all moving actin filaments after removing all filaments whose deviation in velocity is 20% or more than its mean velocity. **(b)** Representative actin-activated ATPase curves for G256E and WT in single-headed sS1 construct showing similar ATPase rates. **(c,d)** Representative actin-activated ATPase curves for WT and G256E two-headed myosin constructs. Each data point represents the average of three technical replicates of one biological replicate (n=3). Actin-activated curves are fitted to Michaelis-Menten kinetics, Error bars represent SD and shaded areas depict the 95% CI of the fits. The two other biological replicates can be found in **Fig S1**. **(c)** Long (25hep) tailed wild-type myosin has a ∼40% reduction in ATPase rate whereas the long tailed G256E myosin has a ∼20% reduction in ATPase rate as compared to the corresponding short (2hep) tailed myosin. **(e)** Single turnover assay depicted as percentage of myosin in SRX state. G256E significantly reduces the fraction of myosin in SRX state, resulting in a 26.5± 2.82% (mean ± SEM) decrease in SRX (N=4). Each data point represents a single experiment, and individual fluorescence decay curves can be found in **Fig S2**. ^∗∗∗∗^ p<0.0001.

### Isolated Myofibrils

Myofibrils isolated from MYH7 G256E mutation-bearing hiPSC-CMs generate increased tension, demonstrating a hypercontractile phenotype.

To examine how dysregulated interactions of G256E mutant myosin with actin affect the contractile properties of myofibrils, we first generated a MYH7 G256E iPSC line (MYH7^WT/G256E^) and the isogenic wild-type counterpart (MYH7^WT/WT^) using CRISPR/Cas9 gene editing technology. WT myofibrils were extracted from isogenic WT hiPSC-CMs (MYH7^WT/WT^), while G256E mutant myofibrils were extracted from MYH7 G256E mutant hiPSC-CMs (MYH7^WT/G256E^), and thus contain both mutant and non-mutant forms of β-cardiac myosin, similar to the human disease. The mechanical and kinetic properties of MYH7^WT/G256E^ and its isogenic control MYH7^WT/WT^ hiPSC-CM myofibrils were measured in a custom-built apparatus using a fast solution switching technique, as previously described^20^. At maximal Ca^2+^ activation (pCa 4.0), MYH7 G256E myofibrils generated ∼2X more force than isogenic controls (**Figure 4a,b**, *p* < 0.001). Furthermore, the rate of activation (*k*_ACT_), which is dependent on crossbridge attachment and cycling rates, was significantly increased for MYH7 G256E myofibrils (**Figure 4c,d**, *p* < 0.05).

**Figure 4.**
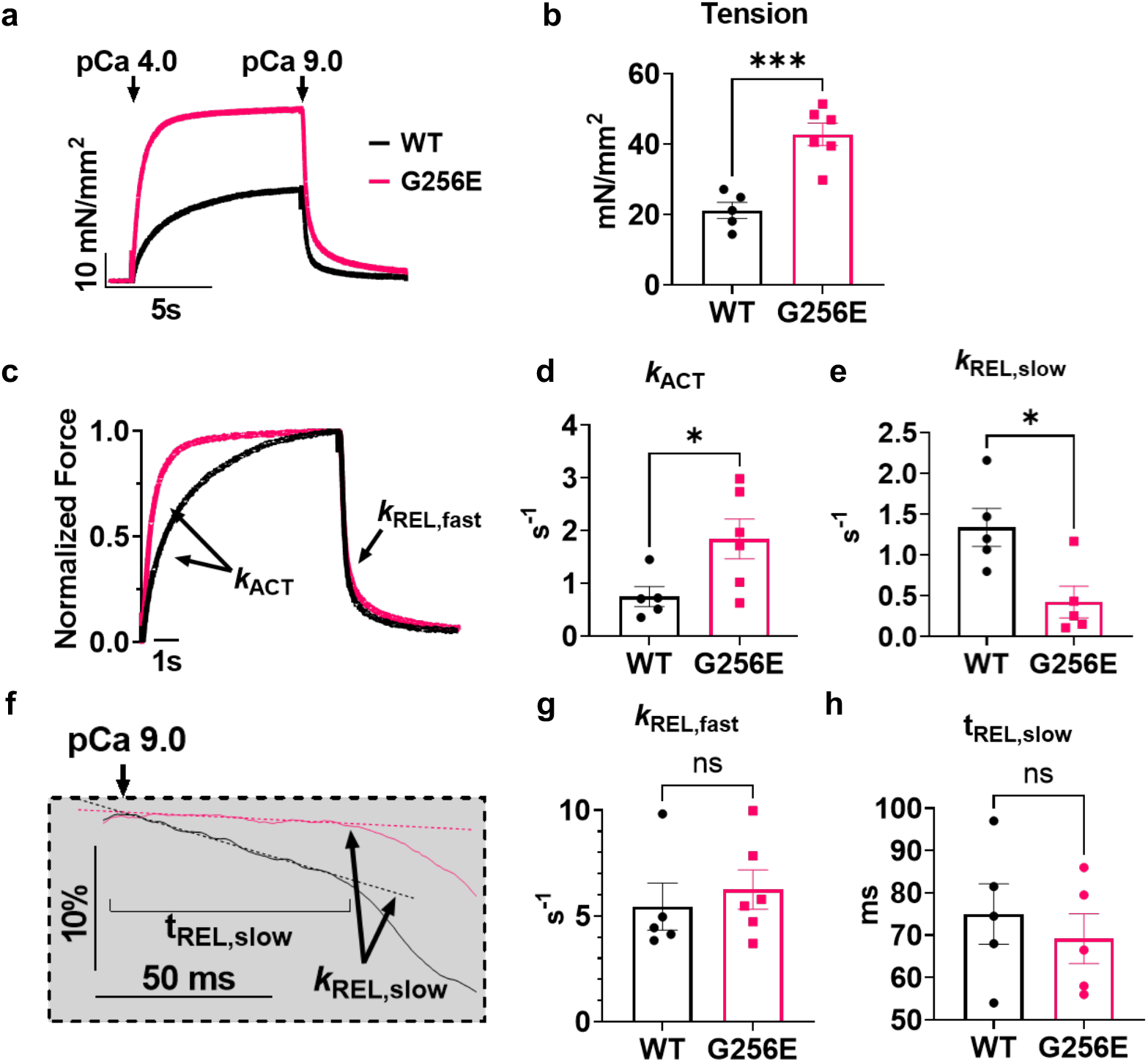
Isolated hiPSC-CM-derived myofibril with G256E mutation demonstrates increased tension generation at a faster rate and delayed slow phase relaxation. **(a,b)** Increased tension can be seen in the activation trace of MYH7 G256E mutant myofibrils (MYH7^WT/G256E^, red) compared to wild-type isogenic controls (MYH7^WT/WT^, black). **(c,d)** Normalized force trace shows increased rate of activation (k_ACT_) of mutant myofibrils compared to isogenic controls. **(e)** Fast phase of relaxation is unaffected. **(f-h)** Close-up of slow phase relaxation showing slower early phase relaxation (k_REL, slow_) of G256E mutants. Statistical significance was determined by Welch’s unpaired two-tailed t-test. ns p>0.05, * p<0.05, *** p<0.001

Relaxation of myofibrils from isometric contraction includes an initial slow, linear phase (*k*_REL,slow_) that is determined by the myosin detachment rate^45^ and a fast exponential phase (*k*_REL,fast_) that reflects multiple active and passive properties of the contractile elements. The *k*_REL,slow_ of MYH7 G256E myofibrils was significantly slower compared to isogenic controls (**Figure 4e,f**, *p* < 0.05) suggesting that the MYH7 G256E mutation slows either ADP release or ATP binding to myosin, the chemo-mechanical steps required for cross-bridge detachment. The duration of the slow, linear phase (t_REL,slow_), which is determined by the rate of thin filament de-activation^46^ and *k*_REL,fast_ were not significantly different between MYH7 G256E myofibrils (**Figure 4g,h**). Overall, the combination of isolated myosin and myofibril mechanics data demonstrate that the MYH7 G256E mutation may result in a hypercontractile phenotype by increasing myosin binding and slowing the rate of cross-bridge cycling to generate greater tension at a faster rate and decrease the slow, early phase of relaxation.

### Single cardiomyocytes

MYH7 G256E mutation-bearing hiPSC-CMs display cellular hallmarks of HCM at the single-cell level.

To examine the effect of the G256E mutation at the next scale level, hiPSC-CMs, we seeded single hiPSC-CMs at 30 days on patterned surfaces with aspect ratios of 7:1 using a 10-kPa hydrogel platform (mimicking physiological morphology and stiffness), which we have shown for optimal force generation^39, 47^, and performed traction force microscopy as previously described^39^. We demonstrated successful patterning and traction force measurement of both MYH7^WT/G256E^ hiPSC-CMs (G256E) and their isogenic controls (MYH7^WT/WT^, WT) (**Figure 5a,b**). Measures from traction force microscopy suggested the G256E mutation resulted in a 2-fold increase in peak traction force generation (***p < 0.0005) and a 1.2-fold increase in contraction velocity (WT: 1.8 µm/s, G256E: 2.2 µm/s), while the relaxation duration (time to 90% relaxation) decreased by 1.2-fold (WT: 0.38s, G256E: 0.3s) (**Figure 5c-e**). Both the WT and G256E hiPSC lines were gene-edited on a line with an EGFP tag at the ɑ-actinin locus (AICS-0075-085), allowing us to monitor sarcomere shortening in real time^48^ (**Figure 5f**). While initial sarcomere length at rest was slightly reduced in the G256E mutants (**Figure 5g**), G256E showed greater sarcomere shortening (WT: 15%, G256E: 20%), indicating increased contraction in G256E mutants at the sarcomere level (**Figure 5h**). Taken together, these data show that G256E mutant hiPSC-CMs show a hypercontractility phenotype at the single cell level.

**Figure 5.**
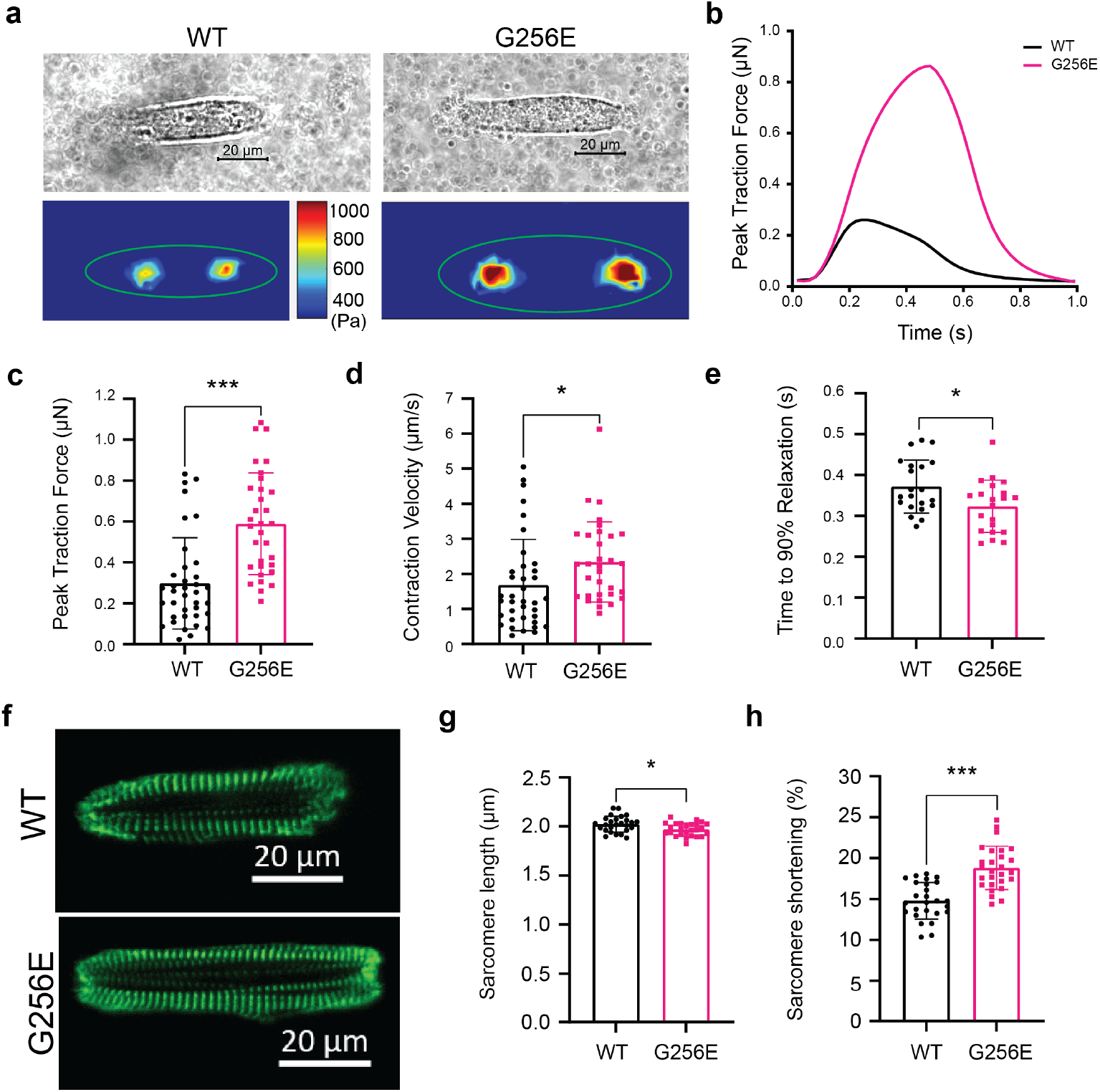
Single cell contractility analysis revealed hiPSC-CMs bearing MYH7 G256E mutation show hypercontractility. (a,b) Representative images of patterned single wild-type isogenic control (MYH7^WT/WT^, WT) and MYH7 G256E mutant hiPSC-CMs (MYH7^WT/G256E^, G256E) and their traction force traces. (c-e) MYH7 G256E mutant (red) depicts increased (c) total peak force, (d) contraction velocity and (e) decreased relaxation velocity compared to wild-type isogenic control. (f) Representative images of sarcomeric organization in single WT isogenic control and G256E mutant hiPSC-CMs. (g, h) The G256E mutant shows statistically reduced sarcomere length and increased sarcomere shortening compared to the WT isogenic control. Data are presented as mean ± standard deviation (SD). Statistical significance was determined by an unpaired t-test. P < 0.05 is designated with (*), P < 0.005 is designated with (**), P < 0.0005 or smaller is designated with (***).

Since cardiac hypertrophy is a clinical hallmark of HCM, we next asked whether the G256E mutation would lead to cellular hypertrophy at the single cell level. We examined cell spread area on 10-kPa hydrogels as well as glass coverslips and found a subtle but significant increase in cell spread area on hydrogel (WT: 800 µm^2^, G256E: 1000 µm^2^; p < 0.005) (**Fig. S3**). In contrast, we did not observe significant changes in cell spread area on glass cover slips of non-physiologic stiffness (WT: 1800 µm^2^, G256E: 1800 µm^2^; p > 0.05) (**Fig. S4**). In sum, G256E mutation-bearing hiPSC-CMs show a robust hypercontractility phenotype in agreement with myosin protein and myofibril data, but less dramatic hypertrophy that is sensitive to substrate stiffness.

### Multicellular engineered heart tissue

The G256E mutation produces a hypercontractile phenotype in multicellular engineered heart tissue.

To examine the effect of the G256E mutation at the multicellular level, three-dimensional engineered heart tissues (EHTs) were generated from WT and G256E hiPSC-CMs. The EHT platform used consists of day 30 hiPSC-CMs and human stromal cells (HS27A) embedded in a fibrin hydrogel and suspended between two elastomeric posts, one flexible and one rigid for longitudinal force measurement^49^. Both WT and G256E hiPSC-CMs produced EHTs that compacted around the posts and initiated synchronized contractions (**Figure 6a**). Tracking the deflection of the flexible post allows for the measurement of contractile force, and G256E EHTs demonstrated an increase in twitch force, specific force, twitch power, and twitch work (**Figure 6b-f**). Analysis of contraction kinetics showed similar times to peak contraction (**Figure 6g**) but an increase in maximum contraction velocity in G256E EHTs as well as shortened times to 50% and 90% relaxation (**Figure 6h-j**). Taken together, these data demonstrate preservation of the hypercontractile phenotype observed at the single molecule, myofibrillar and individual cellular levels to the multicellular level.

**Figure 6.**
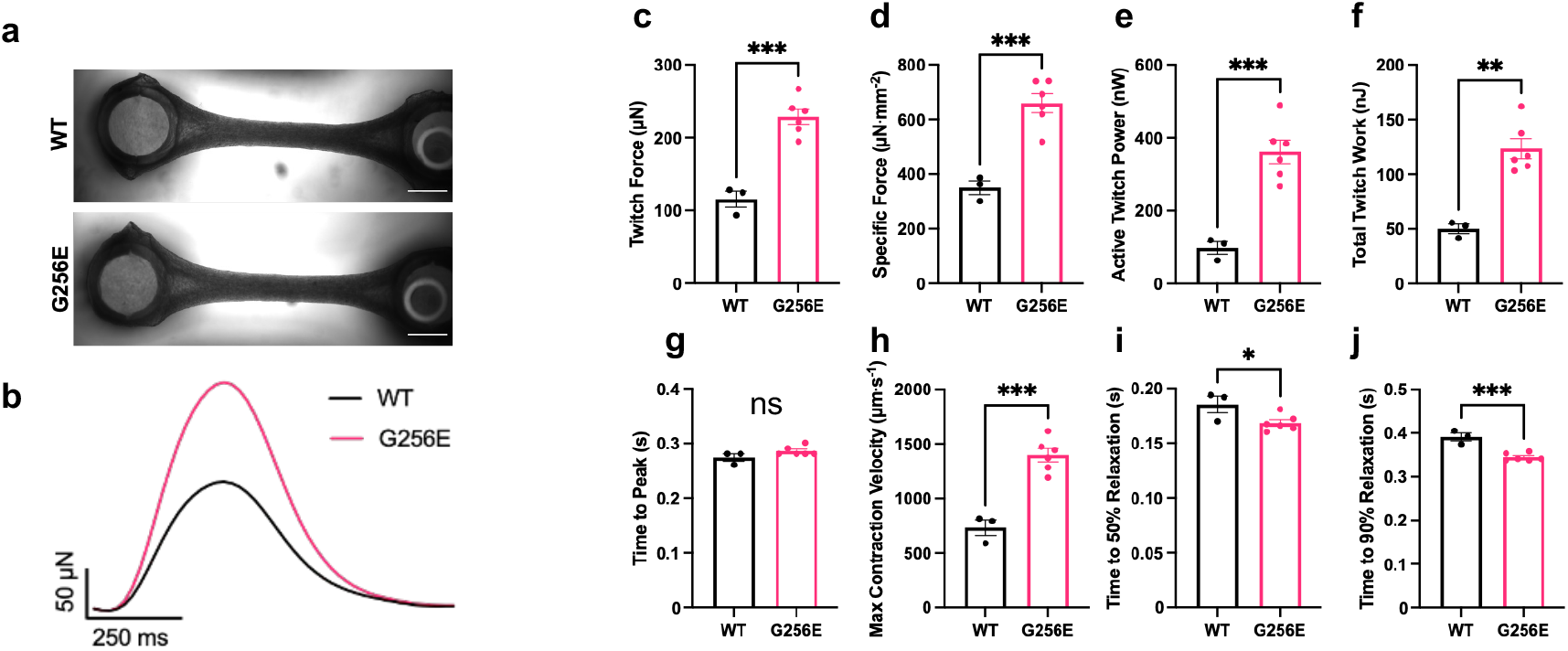
Engineered heart tissue (EHT) model revealed hypercontractility in G256E EHTs. **(a)** Images of representative EHTs, shown with flexible post at the left and rigid post at the right (scale bar = 1 mm). **(b)** Representative traces of EHT twitch force. **(c-j)** Measured metrics of WT and G256E EHT contractility derived from videos of EHT contraction. Each data point represents an individual EHT. * p ≤ 0.05, ** p ≤ 0.01, *** p ≤ 0.001.

Given the central role of calcium handling in modulating cardiac contractility, and prior studies showing a role for altered calcium dynamics in HCM^23, 26^, EHTs were loaded with the Ca^2+^ indicator dye Fluo-4 AM to examine any alterations in Ca^2+^ dynamics that may contribute to the observed contractile phenotype. No significant differences in Ca^2+^ transient kinetics or amplitude were observed between WT and G256E EHTs (**Fig. S5**), indicating that the increased contractility was likely not caused by differences in calcium handling but rather due to direct biomechanical effects at the sarcomere level.

To further investigate potential contributors to the altered contractile kinetics, the relative expression of α-MHC and β-MHC was assessed by immunofluorescence staining. We found that regardless of EHT genotype, most hiPSC-CMs expressed predominantly β-MHC, with only a few, smaller hiPSC-CMs expressing α-MHC (**Fig. S6**). Subcellular sarcomere structure was assessed by immunofluorescence staining for the z-disk protein α-actinin. Both WT and G256E EHTs were shown to have aligned sarcomeres in the direction of EHT contraction, as seen by α-actinin fluorescence, and no difference was observed in either sarcomere length, width of ɑ-actinin layers, or estimated cell area (**Fig. S7**). Taken together, these results further support the G256E mutant myosin’s increased recruitment and altered cycling kinetics.

### Altered cellular metabolism

scRNAseq suggests metabolic alterations in G256E mutation-bearing hiPSC-CMs.

To examine the effect of the G256E mutation on the global transcriptome of hiPSC-CMs, both WT and G256E hiPSC-CMs at day 30 were harvested for droplet-based scRNAseq. The captured cells expressed high levels of cardiomyocyte-specific markers, TNNT2, ACTN2, TNNI1, TNNI3 (**Figure 7a,b**). It is noteworthy that both WT and G256E hiPSC-CMs predominantly expressed MYH7 over MYH6, indicating that day 30 hiPSC-CMs can address the cellular consequences of a MYH7 mutation (**Figure 7b, Fig. S8**). Unbiased clustering revealed 7 sub-clusters of hiPSC-CMs (**Figure 7c**). While Clusters 2 and 1 were composed of a high fraction of G256E mutant cells, 75.9%, 65.2%, respectively, there was no transcriptomically distinct cell cluster, by unsupervised clustering, that distinguishes G256E mutant cells from WT (**Figure 7c,d**).

**Figure 7.**
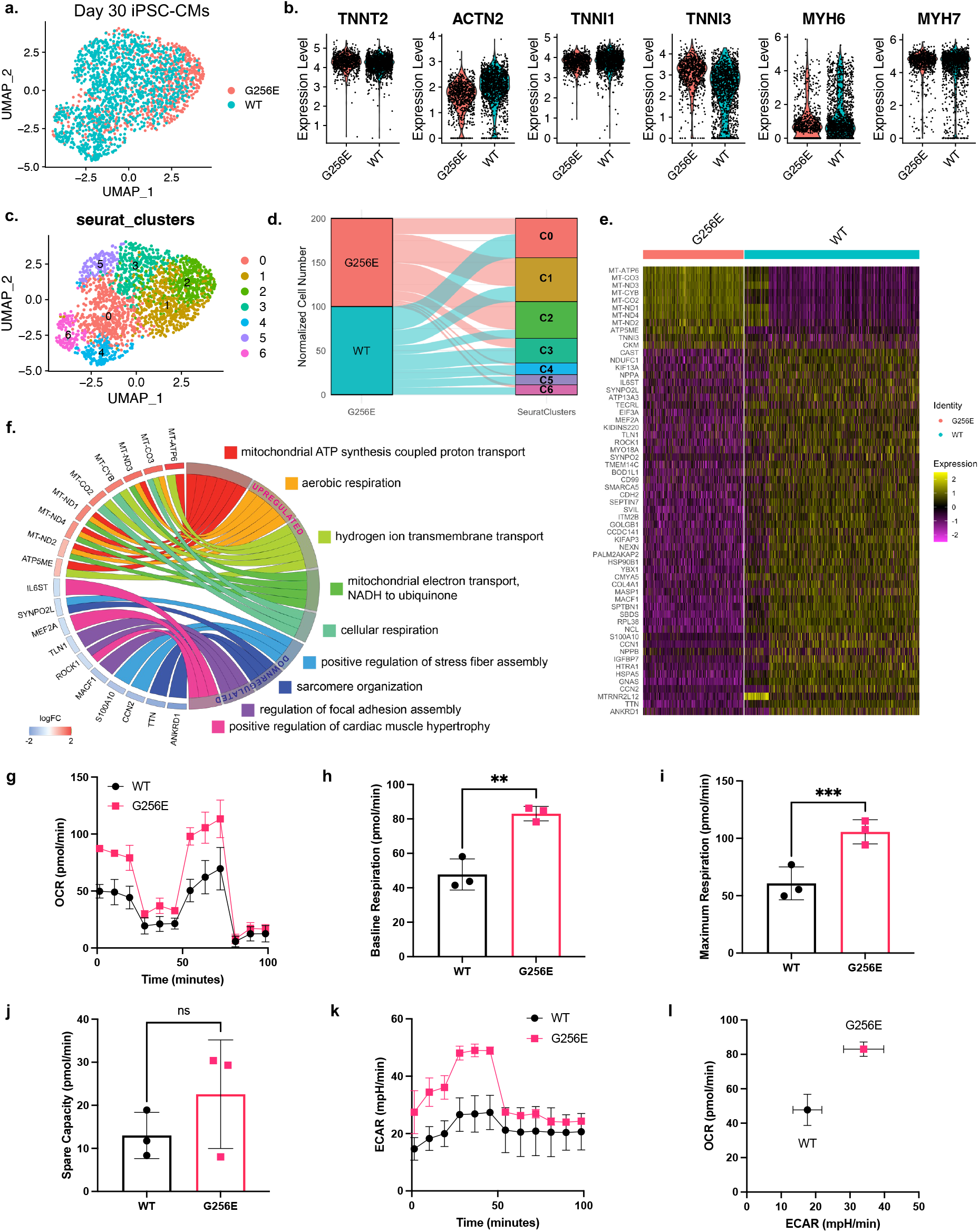
Single cell transcriptomic profiling and Mito Stress test reveal elevated mitochondrial respiration in MYH7-G256E mutant iPSC-CMs. **(a)** UMAP representation of MYH7 G256E mutant cells (MYH7^WT/G256E^, G256E) and the isogenic counterpart (MYH7^WT/WT^, WT) clustered by the presence of G256E mutation. **(b)** Violin plots of cardiomyocyte marker genes. **(c)** Unbiased clustering of G256E and WT iPSC-CMs. **(d)** Alluvial plot demonstrating the composition of Seurat clusters. **(e)** Heatmap of top differentially expressed genes between G256E mutant and isogenic control. (logFC > log1.8). **(f)** Chord plot of gene ontology biological process terms associated with top differentially expressed genes. **(g-l)** Results of mitochondrial respiration profiling. The effect of the MYH7-G256E mutation on **(g)** oxygen consumption rate and **(h)** extracellular acidification rate. **(i)** OCR-ECAR plot demonstrating metabolic shift. **(j-l)** Bar graphs of **(j)** baseline, **(k)** maximum, and **(l)** spare respiration capacity.

Next, we examined the top differentially regulated genes and the associated biological processes in response to the G256E mutation. Top upregulated genes included mitochondrial genes such as MT-ATP6, MT-CO3, MT-ND3 and the corresponding processes they regulate were mitochondrial ATP synthesis coupled proton transport, mitochondrial electron transport, and cellular respiration (**Figure 7e,f, Fig. S9**). On the other hand, traditional hypertrophy marker genes (NPPA, NPPB) and genes reported to be associated with hypertrophic remodeling (SYNPO2L, ANKRD1, CDKN1A, MEF2A) were downregulated (**Fig. S10**), consistent with our finding of a minimal change in cell size.

To further validate the transcriptomic changes related to mitochondrial respiration, we sought to investigate the impact of the G256E mutation on mitochondrial respiratory function using the Seahorse XF Mito Stress test. We found that both baseline and maximal oxygen consumption rate (OCR) of G256E mutant hiPSC-CMs were significantly increased by ∼1.8-fold over WT cells (**Figure 7g-j**), demonstrating increased mitochondrial respiration. Furthermore, we also found elevated extracellular acidification rate (ECAR) in the mutant iPSC-CMs, indicating increased glycolysis (**Figure 7k**). Increase in both OCR and ECAR indicates elevation in oxidative phosphorylation and glycolysis-mediated ATP production (**Figure 7l**). These findings suggest that the hypercontractile G256E mutation increases energy demand in cardiomyocytes, leading to upregulation of mitochondrial genes and increasing mitochondrial respiratory function, as an early feature of HCM.

## DISCUSSION

The multi-scale approach described here allowed us to study the extent to which a specific MYH7 mutation of relatively low penetrance contributes to the primary pathology of hypercontractility and secondary effects of hypertrophy and metabolic remodeling. The MYH7 G256E mutation has been described in a large family and has a relatively low penetrance of hypertrophy of around 50%^34^, leading some to even question its pathogenicity^35^, making it an ideal candidate for assessing its effects using our multi-scale approach. We integrated purified protein assays with a mutation-introduced hiPSC model system on an isogenic background that enabled us to investigate the relationships between biophysical and physiologic mechanisms of HCM, downstream of the G256E mutation. Our primary finding across the multiple scales examined is that the G256E mutation causes hypercontractility at every level.

Molecular dynamics simulations demonstrate that the G256E mutation causes structural changes in the transducer and binding pocket of β-myosin that can directly influence intrinsic sensitivity of the myosin motor to force, reflected through the myosin-actin interaction. This can alter the rate of opening from the folded back (SRX) state and, potentially, multiple steps of the chemo-mechanical crossbridge cycle (**Figure 2**). Myosin motility assays demonstrate that the G256E mutation led to a 20% reduction in actin gliding velocity (**Figure 3a**). Actin gliding velocity is proportional to the step size and force-dependent actin-myosin detachment rate^14, 50^. Our myofibril data demonstrate a 70% reduction in basal detachment rate, *k*_REL,slow_ (**Figure 4e**), which suggests that the mutation may increase the force sensitivity of the actin-myosin detachment rate. Importantly, the G256E mutation led to a decrease in the fraction of myosin in the SRX state, indicating an increase in available open myosin heads that can interact with actin filaments. This effect has been described in other pathogenic MYH7 mutations such as H251N and P710R we previously studied^7, 8, 14^. Increased myosin head availability can contribute to increasing the ratio of ATPase rate of the long-tailed 25-hep myosin to that of the short-tailed 2-hep construct. The ratio_25hep:2hep_ in the G256E mutant was higher than the wild-type myosin indicating hyperactivity, but lower than other myosin mutants known for high penetrance or severe disease phenotype (WT: 0.6, G256E: 1.33, H251N: 1.60, P710R: 1.54) (**Table S1**). The rate of opening from the folded back state (SRX) is also sensitive to force and likely contributes to the increase in max force and rate of force generation observed in the myofibrils. The increase in available myosin heads (recruitment) and slower myosin detachment both increase the number of force-bearing cross-bridges during contractions and, taken together, provide a molecular explanation for the increased tension generation from G256E myofibrils, cells, and multicellular tissue constructs, providing a consistent story across structural and length scales for the G256E mutant hiPSC-CMs (**Figures 5,6**).

Since the transcriptional program orchestrates morphological and functional phenotypic changes of cardiomyocytes, we sought to examine alterations in the transcriptome of hiPSC-CMs induced by the MYH7 G256E mutation. We found that unbiased cell clustering did not lead to separation of G256E mutation-bearing cells from isogenic control cells, indicating an indistinguishable global transcriptome between the two groups (**Figure 7a,c**). MYH7 G256E mutant hiPSC-CMs upregulated genes associated with oxidative phosphorylation such as MT-ATP6, MT-ND1, and MT-ND3 (**Figure 7e,f**). This was corroborated by the Seahorse Mito Stress assay, where MYH7 G256E mutant hiPSC-CMs demonstrated 1.8-fold increased basal and maximal respiration (**Figure 7g-i**). Given increased ATPase activity of double-headed myosins and increased force from the mutant myosin heavy chain (**Figure 3-6**), the increased expression of mitochondrial genes may be an adaptive response to the increased ATP demands of the mutant. Our findings agree with previous HCM studies that have shown altered energetics as proximal consequences of HCM mutations^4, 9, 51, 52^.

Cellular hypertrophy is a hallmark of HCM and is the principal phenotype leading to clinical diagnosis by echocardiography. Hypertrophy is regularly seen in myectomy samples of HCM patients, which often represents a later progression of disease stage, since patients undergoing myectomy tend to have more severe disease. In our hiPSC model, the MYH7 G256E mutation did lead to cellular hypertrophy but only to a modest degree. Cell area measurement of the micro-patterned single-cell on 10-kPa hydrogel did demonstrate a statistically significant increase in cell area in G256E mutant cells (**Fig. S3**). However, this was highly sensitive to substrate stiffness, as G256E mutant cells grown on glass substrates (of non-physiologic stiffness) showed no difference from control (**Fig. S4**).

In addition to cell area, signaling analysis at the gene and protein level also suggested the lack of activation of canonical hypertrophic remodeling, consistent with the minimal hypertrophic phenotype. Previous studies have identified ERK and AKT signaling as essential regulatory signaling pathways for hypertrophic remodeling^14, 53–55^. In our previous study, hiPSC-CMs carrying the early-onset HCM mutation MYH7 P710R were significantly larger than the isogenic control^14^. This enlargement in cell size was found to be linked to enhanced activation of ERK and AKT signaling pathways. In this study, western blot analysis revealed there was no activation of ERK or AKT signaling pathways in G256E mutant cells (**Fig. S11**). This was further supported by scRNAseq data where there were no signs of transcriptomic activation of known hypertrophic remodeling pathways (**Fig. S10**). For example, NPPA, NPPB, well-known hypertrophic markers which have been shown to be upregulated in septal myectomy samples^56^ were downregulated in our study (**Fig. S10a**). P53 signaling activation has been identified as an important regulatory signal that gets activated in response to DNA damage that precedes morphological remodeling in HCM^56^. CDKN1A, the P53 transcriptional target, and HSP90AA1, P53 signaling stress response gene, were both downregulated in G256E hiPSC-CMs (**Fig. S10b**). Another well-known HCM transcriptional regulator that is known to mediate pathological hypertrophic remodeling is myocyte enhancer factor 2 (MEF2)^57, 58^. scRNAseq data indicate MEF2A as one of the top downregulated genes (**Fig. S10c**). Furthermore, Epigenetic Landscape In Silico Deletion Analysis (LISA)^59^ predicted MEF2A as a top transcription factor that orchestrates gene downregulation (P = 3.11 x 10^−9^) that were associated with cardiac muscle hypertrophy (MEF2A, ROCK1, IL6), sarcomere organization (ANKRD1, TTN, SYNPO2L), and focal adhesion assembly (MACF1, TLN1) (**Fig. S10d**). It is possible that other, non-canonical, hypertrophy pathways are activated in mutations like G256E, that show minimal hypertrophic phenotypes *in vitro*. In sum, despite significant increases in myosin recruitment and force in both the G256E and P710R mutations, the data suggests a divergence in hypertrophic response. This indicates that alterations in the functional parameters of myosin are not the sole stimuli for triggering canonical hypertrophic response.

Although a dramatic hypertrophic phenotype was not consistently observed in our hiPSC-CM model, one of the key advantages of using an hiPSC-CM model is the ability to study the pathogenicity of a specific mutation compared to an isogenic control line, reducing the complications of genetic modifiers that would be present in human tissue samples. Future studies are needed to determine how prolonged alterations in mitochondrial gene expression, mitochondrial function and subsequent accumulation of mitochondrial stress will affect HCM disease progression. In addition to this specific MYH7 missense mutation, the effects of additional genetic modifiers as well as environmental factors (such as matrix stiffness, the afterload stress induced by hypertension) need to be addressed in facilitating HCM phenotypic outcome. Finally, since cardiac hypertrophy is usually the metric used for clinical diagnosis of HCM (as opposed to hypercontractility) the disparity of these two phenotypes could in part explain the reported variable penetrance in clinical studies, since patients with HCM are most commonly identified by the degree of left ventricular wall thickening, not by their ejection fraction.

Lastly, a number of limitations of this study should be addressed. The MD simulations were performed on an ATP-bound myosin state (i.e., the post-rigor conformation) for 500 ns; however, myosin undergoes multiple structural transitions between chemo-mechanical states during the crossbridge cycle, which occurs on microsecond to millisecond timescales. The knowledge of structural changes driven by the G256E mutation on the ATP-bound myosin state enabled predictions of which chemo-mechanical states and transitions will most likely be affected by the G256E mutation during the crossbridge cycle. To overcome this limitation, future work can involve MD simulation using starting structures in different states of myosin (e.g., pre-power stroke, actin-bound, etc.). ATPase and SRX/DRX state measurements used shortened myosin constructs that do not form thick filaments *in vitro*, so the potential destabilizing effects of the G256E mutation on SRX heads in the thick filament context was not measured. In single hiPSC-CMs and isolated protein derived from hiPSC-CMs assays, immature protein isoforms (i.e., ɑ-MHC) may still be expressed and contribute to cell-to-cell heterogeneity; however, we did our best to eliminate this factor by directly measuring the β-MHC isoform. Still, differences in the allelic expression of myosin genotypes and myosin isoforms could explain some of the heterogeneity in cell spread area and single cell transcriptomic analysis. Regarding the substrate-dependent effect of cell spread area in response to the G256E mutation, further studies on the impact of substrate stiffness and MYH7 HCM-causing mutations are needed. Another limitation is that by their nature, EHTs contain an averaged phenotype, and heterogenous effects at the cellular level are not captured. In our model system, other tissue structures such as vascular cells and myofibroblasts that are important players in HCM are absent. Understanding the link between altered cellular mechanics and transcriptional changes in the process of cardiomyocyte maturation and adaptation to disease needs to be addressed in the future.

In summary, we demonstrate a collaborative, multi-disciplinary approach to systematically assess the biomechanical effects of an HCM-causing MYH7 mutation, G256E, with incompletely defined clinical pathogenicity, at multiple scales. The use of a multi-scale platform demonstrates the pathogenicity of the G256E mutation, even if it does not lead to all clinical HCM phenotypes *in vitro*. We believe that this unique collaborative multi-scale pipeline provides enhanced insight into the complexities of HCM caused by a MYH7 mutation with uncertain clinical data and provides a paradigm platform for studying other genetic cardiovascular diseases.

## ACKNOWLEDGEMENTS

We acknowledge Dr. Theresia Kraft for a generous gift of an α-myosin antibody ^62^. This work would not have been possible without the Allen Institute for Cell Science team who contributed with cell line generation, especially Rebecca J. Zaunbrecher. The parental wild-type unedited hiPSC line, WTC, was provided by the Bruce R. Conklin Laboratory at the Gladstone Institutes and UCSF. The Allen Institute for Cell Science wishes to thank the Allen Institute for Cell Science founder, Paul G. Allen, for his vision, encouragement, and support.

## AUTHOR CONTRIBUTIONS

Co-first authors (S.L., A.S.V., C.A.B., K.K., S.B.B., M.C.C., D.P.) and co-corresponding authors (D.B., S.M.W., M.R., B.L.P., D.M., K.R., J.S.) conceived the project, designed the experimental pipeline, and took in charge of the different scale evaluation. Specifically, M.C.C. and M.R. designed the *in-silico* experiment for structural evaluation of myosin and conducted molecular dynamics simulations of myosin. D.P., A.S.V., K.R. and J.S. conducted molecular assays and provided significant scientific input in evaluation of mutation effect at the molecular level. K.K., S.M., and M.R. conducted mechanical measurements of myofibril and analyzed/interpreted mechanics and kinetics data of wild-type and mutant myosin myofibrils. C.A.B., A.S.V., O.C., and B.P. conducted single cell phenotyping assays including single cell spread area and sarcomere shortening, and contractility. S.B.B., and D.M. conducted engineered heart tissue phenotyping assays including contractility and Ca^2+^ transient. S.L. and S.M. designed and conducted sample preparation, library preparation, and bioinformatical analyses for single-cell transcriptomic profiling. P.H. and D.L. provided significant assistance with differentiation of human iPSCs, sample/library preparation for 10X Genomics scRNAseq. D.T.P. provided guidance with bioinformatical analyses. S.L., A.S.V., and J.W.J. designed and conducted the experiments for metabolic phenotyping. B.R., J.E.S., and R.N.G. generated iPSC lines used in this study. S.L., A.S.V., C.A.B., K.K., S.B.B., M.C.C., and D.P. wrote the manuscript and all authors contributed to its editing.

## A LIST OF SUPPLEMENTAL MATERIALS

- Extended Methods
- Table S1
- Figures S1 – S11
- References 1 – 28

